# TransAgent: Dynamizing Transcriptional Regulation Analysis via Multi-omics-Aware AI Agent

**DOI:** 10.1101/2025.04.27.650826

**Authors:** Guorui Zhang, Chao Song, Liyuan Liu, Qiuyu Wang, Chunquan Li

**Author notes:** These authors contributed equally: Guorui Zhang, Chao Song, Liyuan Liu.

## Abstract

Transcriptional regulation research, as a core area of life sciences, faces challenges such as scattered multi-omics data, complex joint analysis, and difficulties in integrating data processing tools. To address these issues, we propose TransAgent, an agent software specifically designed for transcriptional regulation analysis. Through innovative designs such as multi-mode operation (planning/execution/automatic), dynamic memory management, rapid MCP tool expansion (integrating over 30 tools), integration of transcriptional regulation annotation data (over 20 data sources including epigenomics and gene expression profiles), and cloud Docker computing, TransAgent significantly improves analysis efficiency. We have successfully applied TransAgent to various transcriptional regulation analysis scenarios such as re-construction of super-enhancer regulatory circuit in esophageal squamous cell carcinoma and identification of key regulators in cardiomyocyte differentiation, demonstrating analytical robustness and uncovering biological insights. TransAgent automates the entire process from raw data processing to advanced analysis, such as joint prediction of multi-omics data, transforming traditionally time-consuming and labor-intensive tasks into a conversation-driven approach. This provides a new paradigm for transcriptional regulation research, centered around large models as the core driver of scalable agent application analysis.

## Introduction

Transcriptional regulation is a complex process that determines when and how different genes work together to control cell growth. Abnormal transcriptional regulation can lead to disease^1^. Analyzing transcriptional regulation not only unravels the mysteries of life activities but also provides new insights for fields such as precision medicine. However, current transcriptional regulation research still faces many challenges, such as the need for personalized analysis due to differential regulatory mechanisms among individuals, and the requirement for complex analysis pipelines to handle large amounts of data, making traditional methods difficult to integrate and cover complex calling relationships. Moreover, since transcriptional regulation involves interactions among multiple omics, researchers need to simultaneously process epigenomic data and transcriptomic data when analyzing regulatory elements like super-enhancers. In-depth transcriptional regulation analysis also requires the use of numerous specialized analysis software, such as ROSE^2^, HOMER^3^, and deepTools^4^. The entire process involves professional data preprocessing (e.g., quality control, standardization), regulatory element annotation, and functional elucidation of results. These steps are not only tedious but may also require manual adjustments at each stage, with high technical barriers for data preprocessing. Additionally, challenges include data format conversion between different tools and the precise selection of various transcriptional regulation software, especially for researchers lacking a professional background in transcriptional regulation analysis. Meanwhile, users often perform dynamic analyses such as genome annotation, gene expression, and target gene identification during the analysis process, and this dynamism may persist throughout the entire workflow. The dynamic combination of analysis steps makes it difficult to standardize and automate the workflow, and the complex calling relationships between tools are hard to manage with existing Agents. Furthermore, the field is populated with numerous functionally dispersed tools, and their intricate input-output relationships further complicate workflow management.

From the initial emergence of GPT-3^5^, which gave large language models the preliminary ability to call tools (such as OpenAI’s WebGPT^6^ and Google’s LaMDA^7^ supporting web search and API calls), to the significant improvement in reasoning capabilities with models like GPT-3.5/4^8,9^, and the emergence of tools like LangChain that enhance Agents’ ability to autonomously plan tasks, the continuous development of agents in recent years, such as MetaGPT^10^, has been notable. Recently, large language models, such as DeepSeek^11^ and ChatGPT^12^, have made significant progress in multiple fields, driving rapid development in life sciences. Among these, various bioinformatics-related Agents have emerged, such as CellAgent^13^, BioAgents^14^, and MRAgent^15^. BioAgents uses large language models (LLM) to handle genomics tasks with performance close to human experts; CellAgent specializes in single-cell data analysis, automatically selecting tools and parameters; and MRAgent can automatically mine disease causality from literature. While these tools have significantly enhanced vertical domain-specific biological analyses through streamlined workflows and simplified tool-calling relationships, they lack flexibility when encountering new data or tasks, and their analysis processes are opaque, with potential security risks. Most critically, none of them can handle the complex calling relationships among various tools in transcriptional regulation analysis or the integration of large-scale epigenomic and expression data. Therefore, we developed TransAgent—a LLM-driven software for transcriptional regulation analysis. Through intelligent task management and flexible tool calling, TransAgent effectively addresses these issues, enabling researchers to complete complex transcriptional regulation tasks more efficiently. To meet different research needs, TransAgent offers three operating modes to adapt to transcriptional regulation analysis scenarios of varying complexity, including Planning Mode: captures user needs precisely through deep interaction, such as transcription factor activity prediction, binding prediction, epigenomic annotation, and gene expression analysis, and can generate detailed analysis workflows to ensure scientific rigor and reproducibility; Execution Mode: flexibly calls various transcriptional regulation tools and provides real-time feedback during execution to ensure stable progress of the analysis workflow; Automatic Mode: completes the entire analysis process without manual intervention, significantly improving data processing efficiency. We applied TransAgent to classic transcriptional regulation analysis tasks such as super-enhancer regulatory circuit re-construction and key regulators identification in cardiomyocyte differentiation, obtaining reliable results as expected.

Additionally, TransAgent can quickly integrate new tools (including custom system-level tools and MCP services^16^) without modifying core code, greatly enhancing scalability. We also use Docker virtualization cloud technology to handle computationally intensive tasks, saving local resources while ensuring speed. For model output standardization, TransAgent requires all outputs to be in JSON format, fully recording each step’s “thinking-tool call-result observation” to ensure traceability and reproducibility. In terms of security, the system prevents user errors or accidental modification of original data through permission control and container isolation technology. TransAgent also excels in dynamic memory management, selectively retaining or hiding past conversation memories to prevent memory loss or confusion due to excessive conversation length or redundant information, which could lead to inaccurate final analysis results. Furthermore, as an Agent specifically designed for transcriptional regulation, it supports zero-code operation, automatically optimizes analysis workflows (e.g., dynamically adjusting peak calling parameters based on ChIP-seq data quality), and flexibly chains tools (e.g., from ChIP-seq differential peak analysis to transcription factor-target gene network construction). It also reuses historical analysis results (e.g., identified super-enhancer regions) through a dynamic memory management system, significantly improving the efficiency of complex tasks. These features make TransAgent an important bridge connecting researchers and computational tools, providing a useful new approach to help researchers solve complex transcriptional regulation problems.

## Results

### TransAgent Software Automates Transcriptional Regulation Analysis Workflow

Transcriptional regulation data sources are complex and analysis methods diverse. When analyzing transcriptional regulation, researchers typically need to manually collect large amounts of specific omics data and perform functional annotation through region annotation methods. However, the variability in complex omics and researchers’ needs for specific tasks makes it impossible to standardize workflows. To address this challenge, we introduce TransAgent, an interactive Agent software focused on transcriptional regulation analysis. TransAgent, with ReAct^17^ as its core architecture, capable of rapidly calling various predefined tools and expanding domain-specific tools through advanced MCP services to meet researchers’ personalized transcriptional regulation analysis needs (**Figure 1A**). Meanwhile, through virtualization container technology, we migrate all computationally intensive tasks to professional-grade servers and execute cloud system-level commands via our software’s remote calling tools, achieving unified rapid local deployment and online analysis (**Figure 1B**). Due to the context length limitations of large language models, Agents exhibit significant “memory loss” in long-interaction tasks. Although several large language models now have context windows of up to 1 million tokens, such as Gemini 2.5 Pro and Qwen2.5-1M^18^, longer contexts are more likely to cause the model to overlook key information. To effectively solve this problem, TransAgent adopts a combination of long-term and short-term memory (**Figure 1C**). Short-term memory retains messages from a few conversation cycles, while long-term memory retains user messages and Agent thinking content several times that of short-term memory. For content the agent needs to recall, we provide a memory backtracking tool to further reduce context length while ensuring key information retention. More critically, our software innovatively introduces dynamic context memory management, allowing users to manually enable or disable interaction history for specific steps, giving TransAgent fine-grained control over analysis workflows and the ability to handle ultra-long context interaction analysis.

**Figure 1.**
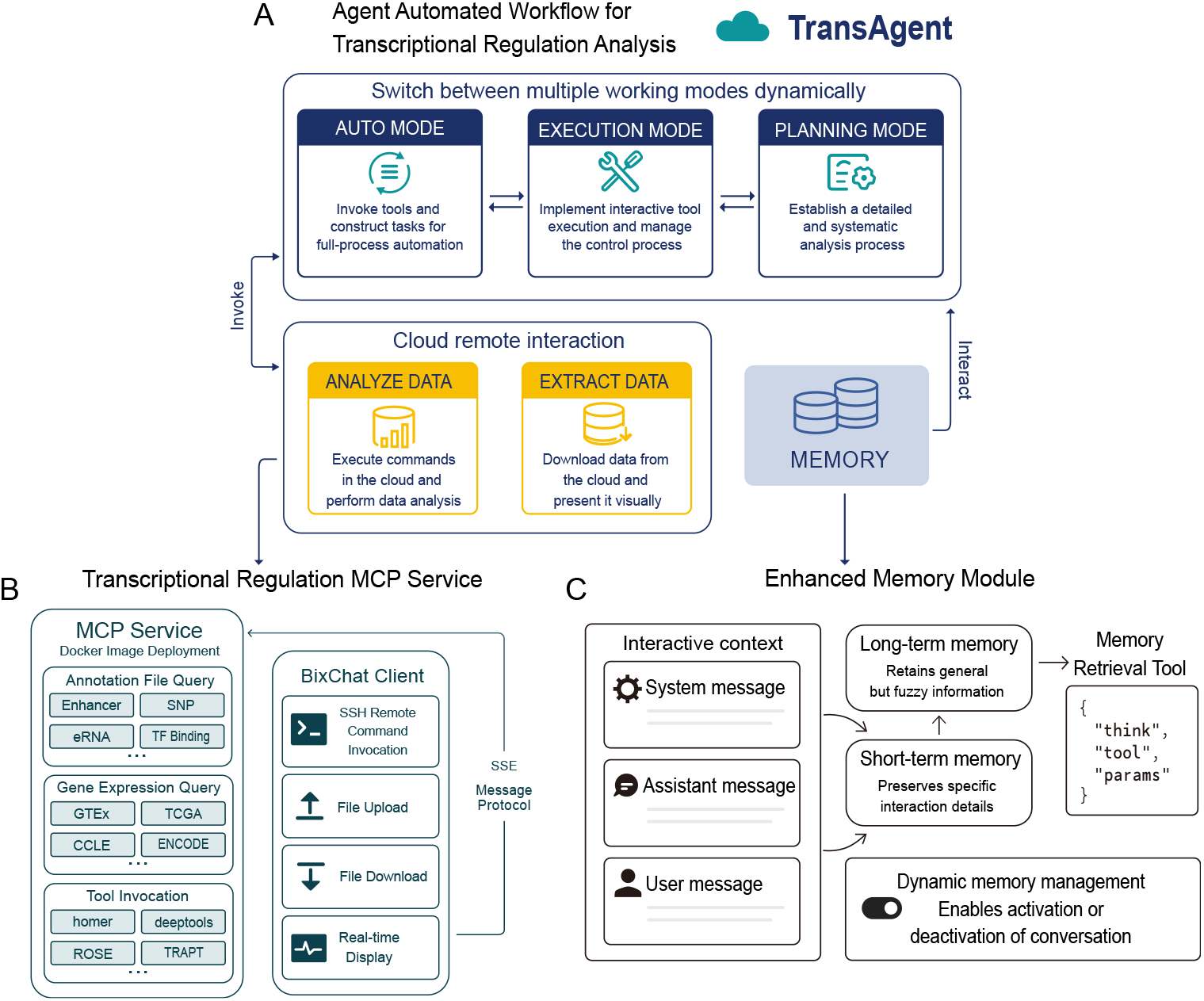
TransAgent Software Architecture. **A** TransAgent core employs three modes: Automatic mode, Execution mode, and Planning mode. Automatic mode calls tools and constructs tasks for full-process automation, Execution mode implements interactive tool execution and manages control flow, while Planning mode is responsible for establishing a detailed and systematic analysis process. Users can dynamically switch between modes to adapt to corresponding tasks. In all modes, commands can be executed in the cloud for data analysis, while data can be downloaded from the cloud for visualization. All interactive context content is saved as memory, enabling real-time operations and interactions. **B** We provide a transcriptional regulation MCP service, where various interactive commands executed through the TransAgent client use the SSE message protocol for bidirectional communication with the cloud MCP Docker environment. In the MCP Docker environment, we offer comprehensive transcriptional regulation annotation data, such as Enhancer, SNP, eRNA, and TF Binding, etc. We also provide various expression data types from multiple sources, such as GTEx, TCGA, CCLE, and ENCODE, etc. Additionally, a large number of transcriptional regulation toolkits are available, such as HOMER, deepTools, ROSE, and TRAPT, etc. **C** For complex transcriptional regulation tasks, TransAgent provides an enhanced memory module with three main types of context messages: system messages, assistant messages, and user messages. In shorter conversation cycles, assistant and user messages are saved as short-term memory, while in longer conversation cycles, most messages retain key information and are saved as long-term memory. TransAgent provides a memory retrieval tool for retrieving detailed information via IDs in long-term memory. For all context memory, TransAgent offers atomic-level memory management, allowing users to precisely enable or disable memory content for specified steps.

TransAgent interacts with the backend large language model through natural language dialogue, with customized interaction modes for different research needs, primarily including three modes: Automatic Mode, Execution Mode, and Planning Mode (**Figure 1A**). Users can switch between these modes to complete various complex transcriptional regulation analysis tasks with maximum controllability. In Planning Mode, TransAgent engages in in-depth interaction with users based on their task requirements, repeatedly asking for more detailed information, including data sources and whether to upload their own data or use local databases. After collecting sufficient information, the Agent provides specific subtasks and then asks the user to manually switch to Execution or Automatic Mode. In Execution Mode, TransAgent automatically thinks and calls appropriate tools based on the current subtask, with the thinking process and tool calling occurring simultaneously. After tool execution, the Agent observes the results and proceeds to the next step of thinking and tool calling until all tasks are completed. Notably, the Agent may encounter failures during execution and will attempt to manually resolve issues. When encountering unsolvable problems, such as missing input files, it will ask the user to upload the corresponding files and pause the current task until user feedback is received, after which the Agent will continue execution. In Automatic Mode, TransAgent is prohibited from interacting with the user, and all encountered problems must be resolved manually, which is a necessary option for fully automated control. Other execution processes remain consistent with Execution Mode until the final task is completed and results are output.

To validate TransAgent’s feasibility in various transcriptional regulation analysis tasks, we used TransAgent to reproduce several classic cases in the field of transcriptional regulation, including region functional annotation, super-enhancer driven regulatory circuit re-construction, and key transcriptional regulator identification. Multiple analysis results demonstrate TransAgent’s general capabilities and broad applicability.

### TransAgent Deciphers Super-Enhancer-Driven Transcriptional Regulatory Circuit in Esophageal Squamous Cell Carcinoma

Super-enhancer-driven transcriptional regulation plays a key role in cancer pathogenesis, and deciphering its molecular mechanisms is crucial for discovering oncogenic drivers. To evaluate TransAgent’s capability in analyzing disease-related regulatory circuits, we investigated the super-enhancer driven regulatory circuit in esophageal squamous cell carcinoma (ESCC). The study first uploaded H3K27ac ChIP-seq raw data (fastq files^19^) to TransAgent, which autonomously initiated the entire analysis pipeline: including data quality control (FastQC), sequence alignment (Bowtie2^20^), and peak detection (MACS2^21^). Through conversational analysis, TransAgent could combine previous analysis results with user prompts to call relevant tools, such as ChIPseeker^22^ software, automatically completing code for analysis and visualization (**Figure 2A**). TransAgent also integrated multiple strategies like geneMapper and BETA^23^ to identify downstream target genes of super-enhancers (**Figure 2B**). Further super-enhancer prediction results from ROSE showed that the identified super-enhancers were highly consistent with previously reported ESCC super-enhancer maps, validating TransAgent’s accuracy in chromatin feature analysis (**Figure 2C**). To deeply decode the regulatory circuit, TransAgent was prompted to use the CRCMapper^24^ tool for analysis, successfully reconstructing the core transcriptional network. By filtering highly expressed genes in ESCC and potential master regulatory transcription factors identified by CRCMapper, we pinpointed 12 key regulators, which also closely matched previously reported results (**Figure 2D**). A correlation network was constructed between these key regulators and ESCC/TCGA ESCA expression profiles (**Figure 2F**), revealing potential relationships among key master regulators like TP63, SOX2, and KLF5—all proven to be important oncogenic transcriptional regulators in ESCC. Meanwhile, genomic binding enrichment analysis of identified peaks using deepTools automatically extracted various local annotation data, revealing significant peak binding at super-enhancer regions, enhancer regions, eRNA regions, and transcription factor binding sites (**Figure 2E**). This further highlights the critical role of epigenetics in maintaining malignant transcriptional programs. This multi-layered analysis not only improved prediction accuracy but also effectively reduced the false-positive rates common in traditional single-method analyses. Notably, TransAgent demonstrated end-to-end automation, analytical robustness, and translational potential through seamless integration of computational tools, making it a powerful platform for accelerating cancer therapeutic target discovery.

**Figure 2.**
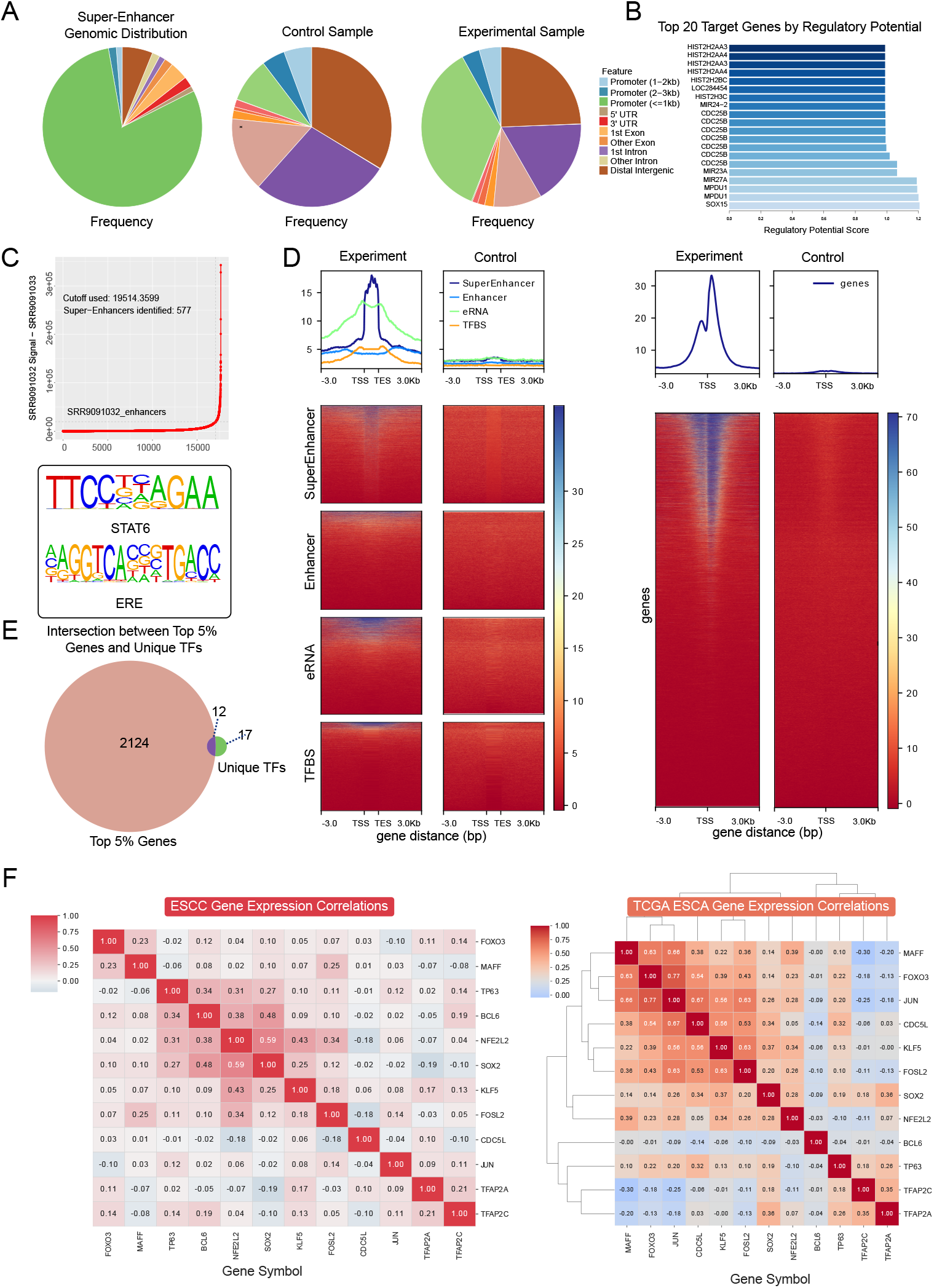
TransAgent Analysis of Super Enhancer-Driven Transcriptional Regulatory Network in Esophageal Squamous Cell Carcinoma. **A** The pie charts from left to right show the genomic proportions of super enhancers identified by ROSE, the control group, and the experimental group. **B** The bar chart displays the top 20 target genes ranked by regulatory potential scores identified by BETA for super enhancers. **C** The upper dot plot shows super enhancers identified by ROSE, while the lower part shows TFs enriched in super enhancer regions. **D** deepTools analysis results, with the left side of each image representing the experimental group and the right side the control group. **E** The upper part of the image shows the overall distribution of various local database region files, including Super Enhancer, Enhancer, eRNA, TFBS, and genes. The lower part shows the coverage of peaks in different region data. **F** The left heatmap shows the expression correlation of key master regulators in ESCC RNA-seq (n=274). The right heatmap shows the expression correlation of key master regulators in TCGA ESCA RNA-seq (n=196).

### TransAgent Identifies Key Transcriptional Regulators and Their Regulatory Networks in Cardiomyocyte Differentiation

Deciphering the transcriptional regulation of cardiomyocyte differentiation is crucial for cardiovascular research. To demonstrate TransAgent’s capability in decoding differentiation-related gene regulatory networks, we studied differentially expressed genes (DEGs) during cardiomyocyte differentiation. The study first input RNA-seq data of the top 200 DEGs from developing cardiomyocytes under two conditions into TransAgent^25–27^. In Planning Mode, TransAgent automatically generated detailed analysis steps (**Figure 3A**). After switching to Execution Mode, TransAgent used the built-in TRAPT^28^ tool to predict upstream transcriptional regulators, successfully screening cardiac master regulators GATA4, NKX2-5, and TBX5, which are known to coordinate heart development through specific transcriptional programs. Following user prompts for genomic task distribution analysis, TransAgent planned the analysis workflow, automatically called ChIPseeker for analysis, and generated visualization code (**Figure 3B**). Here, we observed significant high expression of key transcriptional regulators in cardiac tissue and their binding preferences for promoters and enhancers, confirming the importance of TransAgent-identified regulators in cardiomyocyte differentiation. Additionally, in further target gene prediction tasks, TransAgent constructed a cyclic workflow, iteratively calling the BETA software based on predicted key regulators to infer downstream target genes using regulatory potential (**Figure 3C**). Meanwhile, the gene regulatory network drawn by TransAgent clearly illustrated how these transcription factors collaboratively regulate cardiomyocyte maturation (**Figure 3D**). TransAgent could biologically interpret predicted target genes upon user prompts, further enhancing network credibility, and prioritized high-confidence regulatory interactions, enabling precise analysis and summarization without manual literature mining or computational pipeline setup. Beyond upstream transcription factor identification, we further prompted TransAgent to analyze key target gene expression. TransAgent extracted target gene expression profiles from the GTEx database and automatically generated visualizations (**Figure 3F**). Clustering plots clearly showed enrichment of key targets in cardiac and muscle tissues, further validating TransAgent’s analytical accuracy. To verify the functional significance of the predicted network, we prompted TransAgent to perform enrichment analysis on target gene sets, revealing significant associations with key pathways like cardiac muscle contraction (**Figure 3G**). The system also provided tissue-specific expression profiles of identified regulators (**Figure 3E**), further supporting their roles in cardiomyocyte differentiation. This highlights TransAgent’s value in simplifying regulatory network inference—from DEG analysis to master regulator prediction and functional validation through fully interactive language-driven analysis. The platform automates complex bioinformatics workflows while maintaining biological interpretability, helping researchers efficiently uncover novel transcriptional mechanisms in developmental biology and disease.

**Figure 3.**
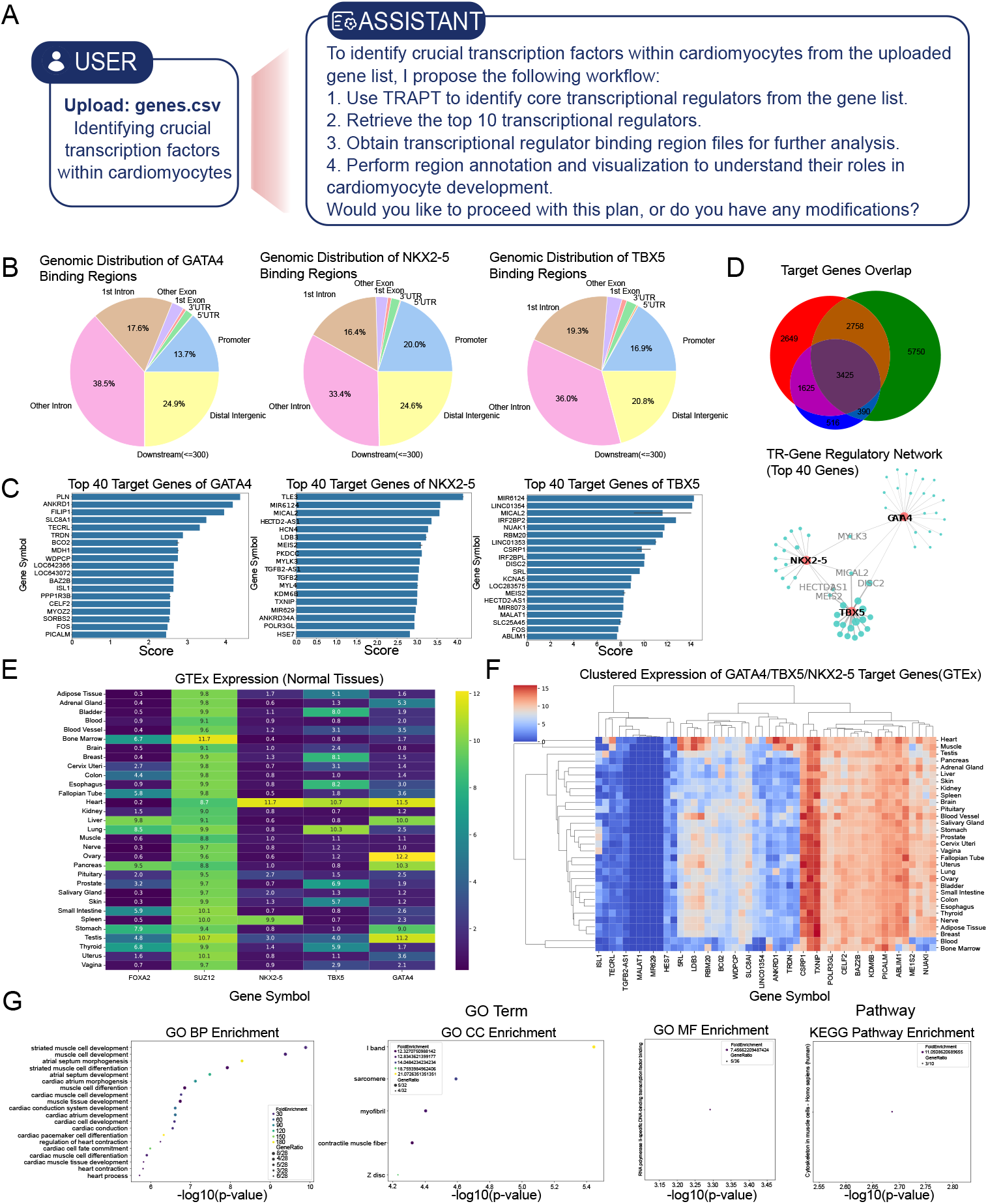
TransAgent Identification of Key Transcriptional Regulators and Their Regulatory Networks in Cardiomyocyte Differentiation. **A** The image shows the data uploaded by researchers and simple prompts, with TransAgent automatically planning detailed execution processes in Planning mode. **B** The pie chart shows the genomic proportions of binding sites identified by ChIP-seq for key transcriptional regulators GATA4, NKX2-5, and TBX5 identified by TRAPT. **C** The bar charts show the average regulatory potential scores of the top 40 non-redundant target genes of key transcriptional regulators identified by BETA software. **D** The upper Venn diagram shows the overlap of high regulatory potential target genes of key transcriptional regulators identified by BETA software. The lower network diagram shows the regulatory network between key transcriptional regulators and high regulatory potential target genes. **E** The heatmap shows the gene expression of key transcriptional regulators in normal human tissues. **F** The clustered heatmap shows the expression of high regulatory potential target genes of key transcriptional regulators in normal human tissues. **G** The bubble plot shows the enrichment of high regulatory potential genes in GO and Pathway terms. The size of the points represents the number of genes covered in the set, and the color represents the enrichment fold.

## Discussion

Transcriptional regulation research is an extremely important direction in life sciences, involving extensive multi-omics and differential data joint analysis. For example, constructing transcriptional regulatory networks requires researchers to manually process complex epigenomic and transcriptomic data and use various transcriptional regulation tools. However, complex data types and extensive tool usage demand advanced data analysis skills and significant time investment with low returns. More critically, researchers spend considerable time adjusting data input/output formats between software and performing repetitive analyses, further reducing overall research efficiency.

To address this, we developed TransAgent, a general-purpose agent software focused on transcriptional regulation research, integrating vast transcriptional regulation datasets and diverse tools. By adopting the ReAct architecture, combining long- and short-term memory, and implementing fine-grained memory flow control, TransAgent can handle highly complex and long-interaction transcriptional regulation tasks. TransAgent is a large language model (LLM)-driven conversational analysis software that excels at understanding human language and automatically formulating detailed analysis workflows based on researcher requirements. Through in-depth interaction, it allows fine-tuning of tasks during early analysis stages. For complex long-interaction tasks, TransAgent supports atomic-level memory control, enabling researchers to precisely manage the entire analysis workflow.

Additionally, TransAgent integrates large-scale epigenomic annotation data, including enhancers, eRNAs, SNPs, and transcriptional regulators, with each data type sourced from multiple databases, resulting in a transcriptional regulation resource library with over dozens of data sources. For transcriptomic data, we collected expression datasets from GTEx^29^, TCGA^30^, CCLE^31^, and ENCODE^32^, covering comprehensive gene data types across human tissues, cancers, and cells. In terms of tools, TransAgent currently integrates dozens of transcriptional regulation tools. Notably, to facilitate rapid tool expansion and custom toolset needs, we equipped TransAgent with MCP service functionality, packaging all tools into a Docker container named “biotools” as an SSE MCP service. Remote MCP access enables TransAgent to rapidly expand capabilities and unify cloud and local operations.

For workflow control, TransAgent introduces three execution modes: Automatic, Execution, and Planning. Researchers can alternate between modes during analysis. Our experiments found that using Planning Mode before or during tasks, followed by Automatic or Execution Mode, yields better results. During interactions, the agent inevitably makes incorrect tool calls or produces erroneous results. Precise memory control, including enabling/disabling memory, dynamically adjusts context length and directs the agent’s focus to user-specified content. This approach is highly effective for LLMs with limited context windows. Furthermore, to retain more interaction history within limited contexts, we combine long- and short-term memory, significantly mitigating model memory loss and context overflow in complex transcriptional regulation tasks. The provided memory retrieval tool allows researchers or the Agent to recall complete tool call information and results stored in memory upon request.

In summary, TransAgent is a cross-platform, extensible, multi-functional transcriptional regulation intelligent software that formulates detailed analysis paths based on user requirements. It allows real-time memory flow control and intervention during analysis, while mode switching enables precise Agent behavior control and enhances planning and execution effectiveness. TransAgent marks a transformative shift in transcriptional regulation analysis from manual to language-interactive automation, not merely improving efficiency but pioneering a new paradigm in the field. With rapid advancements in large language models and expanding context windows, generative AI’s application in transcriptional regulation will further evolve, enhancing TransAgent’s performance. Inevitably, TransAgent’s capabilities are constrained by the LLM’s inherent limitations, such as instruction-following ability, context understanding, and context length. Current cutting-edge Agent research has seen unified models and tool calls, such as the recent GPT-O3^33^ model enabling LLMs with tool execution capabilities. However, no model has achieved this in life sciences. Unifying LLMs with transcriptional regulation tool calls and integrating them with TransAgent, along with expanding TransAgent’s tool set to broaden its applicability, will be a meaningful research direction in transcriptional regulation.

## Methods

### TransAgent Core Architecture

TransAgent maximizes user customization of core modules, including the system prompt module, memory module, tool prompt module, tool definition module, environment information module, model parameter module, etc. In the TransAgent core module, besides customizing tools, it also adapts to external MCP services. In the memory module, we adopted a combination of long-term and short-term memory, effectively extending the model’s memory length and avoiding memory loss issues in long-dialogue tasks. TransAgent provides a universal transcriptional regulation MCP service and uses Docker virtualization and remote system tool calls to enable our software to perform cloud analysis locally.

### System Prompt Module

We employed fixed key system prompts and injectable custom system prompts. Key system prompts include role definition, core tool definition, and mode definition. In role definition, we specified the agent as an all-around AI assistant capable of calling rich tools to complete user tasks, and provided tool call formats and examples. In core tools, we defined multiple tools, including “MCP service call”, “ask user questions”, “plan mode response”, “wait for user feedback”, “memory retrieval”, and “task termination”. In mode definition, we defined three modes: “Automatic mode”, “Execution mode”, and “Planning mode”. In Planning mode, we set the agent’s goal to collect information and obtain context to create a detailed plan to complete the user’s task, and in this mode, we only allow the agent to call the “plan mode response” tool to respond and create the planning process. The user will review and approve the plan, then switch to Execution or Automatic mode to implement the solution. In Execution mode, we specified that the agent can use tools other than “plan mode response” to complete user tasks. In Automatic mode, we specified that the agent does not need to ask the user questions and autonomously completes the subsequent process until the mode changes.

In custom system prompts, we added specific prompts for transcriptional regulation analysis tasks, including key task requirements and precautions. In task requirements, we specified the location where the agent should store analyzed data and prompted the agent to provide users with analyzable data and analysis options. In precautions, we prohibited the agent from modifying core data content and reminded it to pay attention to the formats and differences between MCP and basic tool calls to reduce model hallucinations. In custom system prompts, users can quickly fine-tune the agent’s behavior based on model and task differences. Moreover, TransAgent also provides an option to load custom system prompts with one click.

### Memory Module

The agent’s memory mainly includes three role types: system messages, assistant messages, and user messages. To effectively reduce model hallucinations and solve memory loss issues, we used various methods. To reduce model hallucinations, we standardized the model’s output to be in standard JSON format. For each step of the agent’s output, it must include three parts: “thinking”, “tool”, and “params”. Except for “params”, which may be empty for different tools, the other two are mandatory. In the “thinking” part, the model should provide the current step’s thought process and the reason for tool calls. In the “tool” part, the model should provide the name of the called tool. In the “params” part, the model should provide the parameters needed for the called tool. These collectively form the main body of the current assistant message. The tool execution process and results are also standardized in JSON format and saved as user messages. For system prompts (system messages) and environment details (user messages), they are provided in each model call but are not saved in the message list.

To solve memory loss issues, we defined long-term memory (LTM) and short-term memory (STM) modules. Long-term memory retains longer but vague information, while short-term memory retains shorter but more specific details. More specifically, in long-term memory, we only retain user messages (user task descriptions and called tool names), the “thinking” part of the large model assistant’s content, and indexes of corresponding detailed information. These messages clearly define the current task to be executed, the assistant’s thought process for calling tools, and the names of called tools. Long-term memory is placed at the end of the system prompt as the “memory list” part in each large model request:

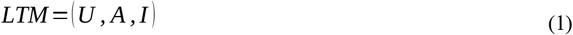

Short-term memory retains all message lists, but during actual model requests, it is truncated based on the user-defined context length, retaining only the most recent interactions:

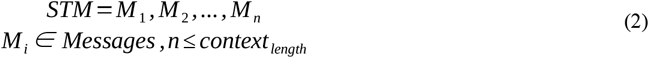

Meanwhile, to enable the agent to recall details from previous interactions, we designed a “memory retrieval” tool. When the agent is asked by the user to recall relevant content or deems it necessary to access detailed information from past interaction message lists, it calls this tool to retrieve key message details based on saved indices.

To further enable TransAgent to perform more complex analyses, we provided precise memory operation functionality. We allow users to dynamically enable or disable interaction history for specific steps during task execution. We offer two options: modular operation and atomic operation, meaning users can choose to enable/disable an entire execution cycle or a single analysis step within a cycle. Notably, this approach allows users to more controllably guide the agent’s behavior path. In ultra-long contexts, users can disable intermediate steps to have the agent analyze initial requirements, such as adding analyses or visualization results. For parallelized processes like repetitive analyses, each cycle can be disabled upon completion, significantly reducing context length.

### Tool Prompt Module

To further enhance the agent’s adjustability and controllability, we allow users to inject core system command prompts for tool calls. Specifically, we provide multiple system command execution tools, including “CLI command execution”, “file display”, “file view”, “file search”, “Python execution”, and “web search”. Among these, “CLI command execution” and “file display” can connect to remote Docker containers via SSH to execute key system commands and tool calls. For rapid tool expansion, we injected definitions and call methods for various transcriptional regulation tools into “CLI command execution”, greatly reducing the complexity of tool definition and providing consistent tool calls, which largely stabilizes the output standardization of large language models by effectively providing few-shot prompts.

### Tool Definition Module

While providing multiple system tools, TransAgent also offers interfaces for user-defined tools. Custom tools follow the same definition format as system tools, including core function definitions and tool prompts. TransAgent also provides several custom tools, such as web search and OCR recognition. This approach allows users to customize their own tools while ensuring they can be called at the same level as system tools, further reducing tool call complexity.

### Environment Information Module

We reserved injection interfaces in both system prompts and environment details. In system prompts, we provide operating system type, platform, and architecture information, which are necessary for the model to correctly execute system commands. For example, when remotely executing Linux commands on a Windows platform, this information must be accurately specified to prevent agent confusion. Environment details are designed to appear after user messages as key environmental information, including current system time, long-term memory lists, and the current language. We reserved an interface for defining the current language to precisely control the model’s response language type.

### Model Parameter Module

The large language model (default DeepSeek model) includes several important parameters, such as context length, temperature, maximum token count, and streaming responses. For instance, the temperature parameter should vary under different circumstances. Lower temperatures yield more stable outputs, which is crucial for the reproducibility of transcriptional regulation analysis tasks. However, this doesn’t mean higher temperatures will cause task failures; they may simply result in different solutions. Different models may have additional definable parameters. For example, in the DeepSeek-Chat model, the response format can be fixed to JSON, which is highly important in our current architectural design. Therefore, we have reserved interfaces for all model parameters to accommodate different tasks and large language models.

### Transcriptional Regulation MCP Service

We provide a universal transcriptional regulation MCP service with multiple defined tools: functional region file queries, gene expression queries, gene coordinate acquisition, bedtools, and system command calls. Within system command calls, we have defined a rich set of transcriptional regulation software packages, such as HOMER, deepTools, ROSE, BETA, TRAPT, and various sequence analysis tools. For functional region file queries, we offer extensive annotation files covering enhancers, super enhancers, SNPs, transcription factor binding, eQTLs, RNA interactions, and CRISPR^34^, with multiple data sources for each type. Gene expression queries also include expression profile data from various sources like GTEx, TCGA, and cell lines (including ENCODE and CCLE).

To simplify deployment and enable high-performance server online analysis, we packaged the MCP service as a unified Docker image using SSE for message transmission. The TransAgent client specifically integrates SSH remote command calls, real-time interactive tools, file upload/download, and display tools. Therefore, TransAgent is recommended as the dedicated client application for such MCP services, offering richer tool execution details and more convenient file transfer/display functions than similar clients.

### Multi-omics annotation data for transcriptional regulation

The TransAgent tool has established a comprehensive annotation system for human genomic regulatory elements, incorporating key database resources independently developed by our research team. For enhancer annotation, the system integrates 2,678,273 super-enhancers from our in-house SEdb^35^ database, along with data from the SEA^36^ and dbSuper^37^ databases. Typical enhancer annotation combines 14,797,266 enhancers from the EnhancerAtlas^38^, HACER^39^, ENCODE, FANTOM5^40^, DENDB^41^, and ENdb^42^ databases, with particular emphasis on 9,176,955 eRNA entries from our proprietary eRNAbase^43^ database.

Genetic variant annotation comprises 37,302,978 common SNPs from dbSNP^44^ (all with minor allele frequency >0.05), 351,728 GWAS risk SNPs from GWAS Catalog^45^ and GWASdb^46^, and 11,995,221 eQTL loci from GTEx, PancanQTL^47^, seeQTL^48^, SCAN^49^, and Oncobase^50^ databases. Chromatin accessibility annotation includes over 130,000,000 ATAC-seq open regions from our ATACdb^51^ database and 69,860,705 DNase hypersensitive sites from ENCODE. 3D genome data incorporates 34,342,926 chromatin interaction sites from 4DGenome^52^ and Oncobase^50^ databases, along with 72,019 TAD structures from 3D Genome Browser^52^. DNA methylation data covers 166,855,665 whole-genome bisulfite sequencing sites and 30,392,523 450K array sites from ENCODE.

Furthermore, TransAgent employs BETA and geneMapper algorithms to effectively identify associations between DNA regulatory elements and target genes. By integrating gene expression profiles from GTEx, TCGA, ENCODE, and CCLE databases, the tool optimizes target gene selection for regulatory elements, providing a comprehensive solution for transcriptional regulation studies.

## Code Availability

The TransAgent implementation code and software described in this article are open-source. See the official TransAgent GitHub repository: https://github.com/TOSTRING-Z/transagent

## Acknowledgments

This work was supported by the Science and Technology Innovation Program of Hunan Province [2024RC1062, 2024RC3212]; National Natural Science Foundation of China [62171166, 62302206, 62031003]; Research Foundation of the First Affiliated Hospital of University of South China for Advanced Talents [20210002-1005 USCAT-2021-01]; Provincial Key Laboratory of Multi-omics and Artificial Intelligence of Cardiovascular Diseases [2023TP1047]; Natural Science Foundation of Hunan Province [2025JJ50401, 2023JJ40594, 2023JJ30536]; Clinical Research 4310 Program of the University of South China [No. 20224310NHYCG05].

